# Learning association for single-cell transcriptomics by integrating profiling of gene expression and alternative polyadenylation

**DOI:** 10.1101/2021.01.04.425335

**Authors:** Guoli Ji, Wujing Xuan, Yibo Zhuang, Lishan Ye, Sheng Zhu, Wenbin Ye, Xi Wang, Xiaohui Wu

**Author notes:** **Guoli Ji** is a professor with the Department of Automation in Xiamen University. His research interests include bioinformatics, advanced control, data mining and information system. **Wujing Xuan** is a graduate student with the Department of Automation in Xiamen University. His research interests are bioinformatics and data mining. **Yibo Zhuang** is an employee in Xiamen YLZ Yihui Technology company. His research interests are software design, cloud computing and big data. **Lishan Ye** is the director of Xiamen Health and Medical Big Data Center. Her research interests are cloud computing and healthcare big data. **Sheng Zhu** is a Ph.D. candidate with the Department of Automation in Xiamen University. His research interests are bioinformatics and healthcare big data. **Wenbin Ye** is a Ph.D. candidate with the Department of Automation in Xiamen University. Her research interests are bioinformatics and mRNA processing. **Xi Wang** is a graduate student with the Department of Automation in Xiamen University. Her research interests are bioinformatics and data mining. **Xiaohui Wu** is an associate professor with the Department of Automation in Xiamen University. Her research interests are mRNA processing, bioinformatics, and data mining. Corresponding author., Tel: 86+13459029440.

## Abstract

Single-cell RNA-sequencing (scRNA-seq) has enabled transcriptome-wide profiling of gene expressions in individual cells. A myriad of computational methods have been proposed to learn cell-cell similarities and/or cluster cells, however, high variability and dropout rate inherent in scRNA-seq confounds reliable quantification of cell-cell associations based on the gene expression profile alone. Lately bioinformatics studies have emerged to capture key transcriptome information on alternative polyadenylation (APA) from standard scRNA-seq and revealed APA dynamics among cell types, suggesting the possibility of discerning cell identities with the APA profile. Complementary information at both layers of APA isoforms and genes creates great potential to develop cost-efficient approaches to dissect cell types based on multiple modalities derived from existing scRNA-seq data without changing experimental technologies. We proposed a toolkit called scLAPA for **<u>l</u>**earning association for **<u>s</u>**ingle-**<u>c</u>**ell transcriptomics by combing single-cell profiling of gene expression and **<u>a</u>**lternative **<u>p</u>**oly**<u>a</u>**denylation derived from the same scRNA-seq data. We compared scLAPA with seven similarity metrics and five clustering methods using diverse scRNA-seq datasets. Comparative results showed that scLAPA is more effective and robust for learning cell-cell similarities and clustering cell types than competing methods. Moreover, with scLAPA we found two hidden subpopulations of peripheral blood mononuclear cells that were undetectable using the gene expression data alone. As a comprehensive toolkit, scLAPA provides a unique strategy to learn cell-cell associations, improve cell type clustering and discover novel cell types by augmentation of gene expression profiles with polyadenylation information, which can be incorporated in most existing scRNA-seq pipelines. scLAPA is available at https://github.com/BMILAB/scLAPA.

## Introduction

Single-cell RNA-sequencing (scRNA-seq) has enabled transcriptome-wide profiling of gene expressions in individual cells, which has great potential to reveal cellular composition of tissues, transcriptional heterogeneity among cells and structure of cell types [1]. Cell-type identification is a critical step in most scRNA-seq data analyses, and a myriad of computational methods have emerged to detect novel cell types, previously un-appreciated sub-types of cells and rare cells [2]. Fundamentally, these numerous clustering methods rely on cell-cell associations (or similarities) for categorizing individual cells into different clusters [3]. A wide range of computational tools have been proposed to cluster cells, which implicitly or explicitly rely on a similarity concept [2]. SIMLR (Single-cell Interpretation via Multikernel Learning) adapts k-means by simultaneously training a similarity measure based on multiple kernel learning [4]. RaceID extends k-means with outlier detection to discover rare cell types [5]. SC3 (Single-Cell Consensus Clustering) utilizes a consensus approach to combine multiple clustering solutions [6]. PhenoGraph combines shared nearest-neighbour graphs and Louvain community detection to fast identify cell clusters [7]. Despite of the considerable progress, there is no strong consensus on which is the best clustering approach to define cell types for all situations [2, 8, 9]. Particularly, high variability and dropout rate inherent in scRNA-seq confounds the reliable quantification of lowly and/or moderately expressed genes [10, 11], resulting in extremely sparse gene-cell count matrix. Consequently, there might be little satisfactory overlap of observed genes among cells, hindering reliable quantification of cell-cell similarities based on the gene expression profile alone.

Recently, multi-omics methods that leverage additional aspects of the cell, such as the DNA methylome, open chromatin or proteome, are beginning to appear [12]. Seurat v3 [13] harmonizes scRNA-seq and scATAC-seq data from a similar tissue to identify subpopulations of cells that are undistinguishable using the scATAC-seq data alone. LIGER [14], a method based on integrative non-negative matrix factorization (iNMF), was proposed to classify cortical cells profiled from single-cell bisulfite sequencing by integrating scRNA-seq data. Additional modalities of individual cells provide valuable information about the phenotype and genetic cellular state not manifested by the transcriptome. However, not all scRNA-seq data is accompanied data from different modalities. Even that multimodal omics data are gradually available, integrative multimodal analysis is still in its infancy [12]. It remains a challenge to reconcile the heterogeneity across modalities as different modalities are normally profiled from cells sampled from the same tissue rather than the same cells.

Although most scRNA-seq studies focus on gene expression profiling, key information on transcript isoforms, e.g., alternative splicing (AS) and/or alternative polyadenylation (APA), can be obtained, enabling multiple aspects of transcriptome information to be derived from standard scRNA-seq without changing experimental technologies [15–20]. Lately, several computational methods, such as scAPAtrap [15], Sierra [16] and scAPA [18], have been proposed to identify APA sites in single cells from diverse 3’ tag-based scRNA-seq protocols, e.g., Drop-seq [21], CEL-seq [22] and 10x Genomics [23]. Cell-to-cell heterogeneity in APA site usage was also observed [15–18]. Particularly, the previous study [15] revealed that the APA profile, even that from non-differentially expressed genes, can distinguish mouse cells in different stages during sperm cell differentiation, suggesting the possibility of discerning cell identities with APA usages independent of gene expression. Recent efforts have pioneered methods to identify APA sites or explore APA dynamics across different cell types [16–18, 24–26], however, most studies profiled APA among cells with predefined cell type labels rather than discern cell types in an unsupervised manner. Complementary information at both layers of APA isoforms and genes can be refined from the same cells [15–20], which creates great potential to develop more sophisticated and cost-efficient approaches to dissect cell types based on multiple modalities derived from existing scRNA-seq experiments.

Here we proposed a toolkit called scLAPA for **<u>l</u>**earning association for **<u>s</u>**ingle-**<u>c</u>**ell transcriptomics by combing single-cell profiling of gene expression and **<u>a</u>**lternative **<u>p</u>**oly**<u>a</u>**denylation. scLAPA leverages the resolution and huge abundance of scRNA-seq, boosting the gene-level analysis with additional layer of APA information directly derived from the same scRNA-seq data. By employing the strategy of similarity network fusion, scLAPA effectively learns highly informative cell-cell associations from expression profiles of both genes and APA isoforms. We compared scLAPA with seven similarity metrics and five clustering methods, using diverse scRNA-seq data from different experimental technologies and species. Comparative results showed that scLAPA is more effective and robust for learning cell-cell similarities and clustering cell types than competing methods. Moreover, with scLAPA we found two hidden subpopulations of cells in peripheral blood mononuclear cells (PBMCs) that were undetectable using the gene expression data alone. As a comprehensive toolkit, scLAPA provides a unique strategy to learn cell-cell associations, improve cell type clustering and discover novel cell types by augmentation of gene expression profiles with polyadenylation information, which can be incorporated in many other standard scRNA-seq pipelines for single-cell analyses.

## Materials and methods

### scRNA-seq datasets

We used five publicly available scRNA-seq datasets from animals and plants generated by 3’ tag-based scRNA-seq protocols (Table S1), spanning a wide spectrum of tissues, cell types and species. Raw data except for the PBMC data were downloaded from NCBI GEO (Gene Expression Omnibus). Cell types and cell labels of the data of Amygdala, Mammary and Root were obtained from the corresponding studies; cell labels of the Hypothalamus data were obtained from PanglaoDB [27]. The PBMC 4k dataset was downloaded from the 10x Genomics website (https://www.10xgenomics.com/). For cell type annotation of PBMCs, we followed the tutorial of Seurat v3 [13] to cluster cells on the basis of the gene-cell expression matrix. Specifically, cells with total read counts less then 300 were discarded. The *LogNormalize* method was adopted for normalization. Top 2000 highly variable features were selected by the *vst* method. PCA (Principal Component Analysis) was used for dimensionality reduction and top 20 principal components were retained. Finally, cells were clustered by Seurat’s *FundClusters* with argument ‘resolution=0.9. For cell type annotation of cell clusters, known marker genes of PBMCs were complied from relevant studies (Table S2). Differentially expressed (DE) genes for each cell group were calculated with Seurat’s *FindAllMarkers*. We also calculated, for each cell cluster, the number of cells where a DE gene is expressed and the mean expression level of a DE gene. The cell type was carefully assigned to a cell cluster according to the presence and expression level of marker gene(s).

### Overview of scLAPA

scLAPA mainly consists of four modules (Figure S1): (i) the input module, (ii) cell-cell distance, (iii) distance fusion, (iv) cell type clustering. The input module prepares the input for scLAPA, including a poly(A) site expression matrix (hereinafter referred as *PA-matrix)* and a gene expression matrix (hereinafter referred as *GE-matrix).* The *PA-matrix* is generated from raw scRNA-seq with scAPAtrap [15], which stores expression levels of poly(A) sites, with each row denoting a poly(A) site and each column denoting a cell. The *GE-matrix* can be obtained from websites like NCBI GEO and 10x Genomics, or generated by various routine scRNA-seq analysis tools like Cell Ranger. In the module of cell-cell distance, a cell-cell distance matrix is learned for *PA-matrix* (called *PA-dist)* and *GE-matrix* (called *GE-dist),* respectively. The module of distance fusion employs similarity network fusion (SNF) [28] to integrate the two distance matrices *(PA-dist* and *GE-dist)* into one cell-cell distance matrix. The cell type clustering module clusters cells based on the fused distance matrix with various clustering methods. scLAPA was implemented as an open source R package and is available at https://github.com/BMILAB/scLAPA. Scripts and data used in this study are also available at the GitHub website.

### Identification of poly(A) sites from scRNA-seq

We followed the tutorial provided at the scAPAtrap website (https://github.com/BMILAB/scAPAtrap) to identify poly(A) sites with scAPAtrap [15]. It should be noted that alternative tools, such as Sierra [16] and scAPA [18], can also be used. Briefly, raw FASTQ reads were mapped with Cell Ranger 2.1.0 (https://www.10xgenomics.com/) and then uniquely mapped reads were obtained with SAMtools (http://samtools.sourceforge.net/). Then UMI-tools [29] was employed to remove polymerase chain reaction (PCR) duplicates and extract unique molecular identifiers (UMIs). The *findTails* function in the scAPAtrap package was used to determine exact locations of poly(A) sites from reads with A/T stretches and the *findPeaks* function was adopted to identify all potential peaks of poly(A) sites from the whole genome level. Finally consensus poly(A) sites supported by both of the peak and the tail evidence were used. The *featureCounts* function in the Subread toolkit [30] was adopted to quantify the expression level for each poly(A) site.

Poly(A) site annotation was performed with the movAPA package [31], using the latest genome annotation of the respective species -- TAIR10 for Arabidopsis, mm10 for mouse and GRCh38 for human. Briefly, poly(A) sites identified from scAPAtrap were annotated with rich information, such as genomic regions (i.e., 5’ UTR, 3’ UTR, coding sequence (CDS), intron, exon and intergenic) and gene id. Similar to previous studies [32–35], annotated 3’ UTRs were extended by a length of 1000 bp to recruit intergenic sites that may originate from authentic 3’ UTRs.

### Calculation of cell-cell distance

scLAPA learns a cell-cell distance matrix for *PA-matrix* and *GE-matrix,* respectively. Various distance metrics can be chosen, including Euclidean distance, Pearson correlation, two metrics of proportionality *(ρ_p_* and *Ø_s_*) [3], RAFSIL (RAndom Forest based SImilarity Learning) [36] and SIMLR [4]. Euclidean distance and Pearson correlation are widely used in either single-cell or bulk transcriptomics. The two measures of proportionality were found to have strong performance according to a comprehensive benchmarking analysis of a large single-cell transcriptome compendium [3]. RAFSIL is a random forest based approach that learns cell-cell similarities from scRNA-seq data, including two variations -- RAFSIL1/2. SIMLR learns a distance metric that fits the structure of the scRNA-seq data by combining multiple kernels corresponding to different informative representations of the data. Euclidean distance and Pearson correlation were calculated by the *dists* and *cor* functions in the R package stats, respectively; SIMLR metric was calculated by the SIMLR R package with argument ‘cores.ratio=0’; RAFSIL metric was calculated by the RAFSIL R package with arguments ‘nrep=50, gene_filter=FALSE’; *ρ_p_* and *Ø_s_* were calculated by the *perb* and *phis* functions in the R package propr, respectively. For each distance metric, cell-cell distance matrices, *PA-dist and GE-dist,* can be learned for *PA-matrix* and *GE-matrix,* respectively. *PA-dist* represents the cell-cell similarity network learned from the APA isoform layer, whereas *GE-dist* reflects the network learned from the gene layer, each of which encapsulates complementary information about cell-cell associations absent in the other genomic layer.

### Distance fusion

After learning *PA-dist and GE-dist,* similarity network fusion (SNF) [28] is utilized to flexibly integrate the two layers of cell-cell similarities into one similarity matrix. First, *PA-dist and GE-dist* were iteratively and gradually fused to a consensus network, utilizing the non-linear method of message passing theory [37]. Then weak similarities representing potential noise were discarded, and strong similarities were retained. By generating coherent cell-cell similarities from both APA isoform and gene layers, SNF profiles a more comprehensive biological relationship among cells, beyond the scope of methods solely based on *GE-matrix.*

Given a *PA-matrix* storing expression levels of *m* poly(A) sites in *n* cells or a *GE-matrix* recording expression levels of *m* genes in *n* cells, the corresponding cell-cell distance matrix *(PA-dist* or *GE-dist*) can be obtained using a selected distance metric. The distance matrix can also be denoted as a graph *G* = < *V, E, W* >, with vertices *V* {*c*_1_, …, *c_n_*} corresponding to cells, edges *E* representing cell-cell link and edge weights *W*_[*n×n*]_ denoting the kernel representation of cell-cell similarities. The weight of an edge linking cells *c_i_* and *c_j_* is determined using a scaled exponential similarity kernel:

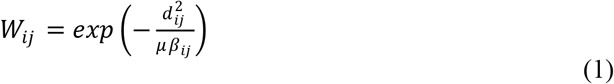

Here *d_ij_* represents the distance between cells *c_i_*, and *c_j_* measured by a distance metric (e.g. Pearson correlation). *μ* is an empirical hyperparameter with a recommended value in a sizable range of [0.3, 0.8] [28]. *β_ij_* is a scaling factor defined as follows:

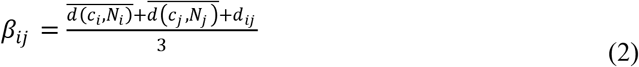

where *N_i_* are neighboring cells of *c_i_* and 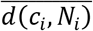 is the average distance of *c_i_* to its neighbors.

To obtain a fused network from *PA-dist* and *GE-dist,* a full and sparse kernel on the vertex set *V* is derived from the weight matrix *W*. The full kernel is a normalized weight matrix 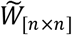 which stores the full information of cell-cell similarities. The normalized weight between *c_i_*, and *C_j_* is defined as:

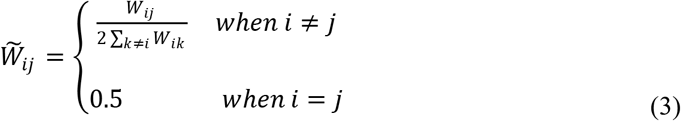

Another matrix *A*_[*n×n*]_ encodes the local affinity that measures similarities of a cell to its *K* most similar cells:

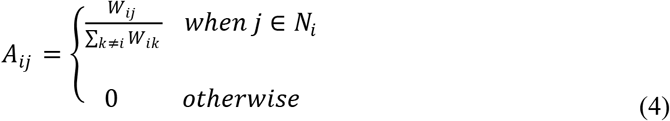

Here *N_i_* is the set of cell *c_i_* and its neighbors in the graph *G.* The network fusion initiates from 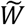, using *A* as the kernel matrix to capture local structure of the graph.

To fuse the two distance matrices *(PA-dist* and *GE-dist)*, first *W^PA^* and *W^GE^* were computed, respectively. Then the corresponding initial state matrices 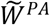 and 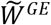 were derived from the two similarity matrices, and the kernel matrices *A^PA^* and *A^GE^* were also computed. Given the initial two status matrices at 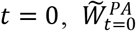 and 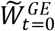, the fusion process iteratively updates the respective similarity matrix:

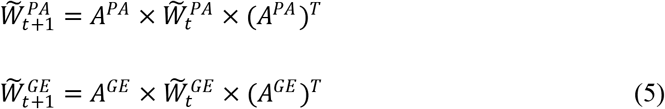

Then after *t* iterations, the final status matrix is obtained:

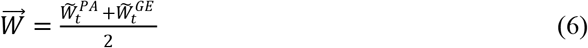

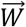 is the fused cell-cell distance network by incorporating cells’ APA isoform and gene expression profiles. The corresponding cell-cell similarity matrix is 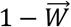. The distance or similarity matrix can be used for downstream cell type clustering.

### Single cell clustering

Four widely-used clustering methods were provided in scLAPA to cluster cells on the basis of the fused cell-cell similarity matrix, including Louvain clustering [38], hierarchical clustering (HC) [39], spectral clustering (SC) [40] and k-means. The Louvain clustering was implemented by the *cluster_louvain* function in the R package igraph, with arguments ‘mode=undirected, weighted=TRUE, diag = TRUE’. The spectral clustering was implemented by the *SpectralClustering* function in the R package SNFtool with default settings [28]. The hierarchical clustering [39] was performed by the *flashClust* function in the R package flashClust with default settings [41]. The k-means clustering was implemented by the *kmeans* function of the R package stats with arguments *‘*iter.max= 1e+9, nstart= 1000’.

### Performance evaluation

We distinguished two scenarios, similarity learning and clustering, to evaluate our approach. For each scenario, we applied scLAPA to four scRNA-seq datasets with pre-annotated cell labels, and compared results with other competing approaches. For the scenario of similarity learning, we compared scLAPA with seven similarity measures, including three measures designed for scRNA-seq (RAFSIL1/2 and SIMLR), two measures of proportionality *(ρ_p_* and *Ø_s_*) and two traditional similarity measures (Euclidean distance and Pearson correlation). Each of these measures was applied to a given *GE-matrix* to learn a cell-cell similarity matrix. For scLAPA, we applied each measure to learn two cell-cell similarity matrices from *PA-matrix* and *GE-matrix* and fused them into one matrix. We also applied different clustering methods including Louvain, HC, SC and k-means on the similarity matrix learned from each similarity measure to assess different similarity measures in the context of clustering.

For the scenario of clustering, we compared scLAPA with five state of the art clustering methods for scRNA-seq data, including SC3 [6], Seurat v3 [13], SINCERA [42], SNN-Cliq [43] and dynamic tree cut method (dynamicTreeCut) [44]. None of these approaches provides explicit similarity learning procedure, instead they provide cell labels by unsupervised learning on the *GE-matrix*. Each approach was applied to a given *GE-matrix* for cell clustering and class labels of cells were obtained. For scLAPA, we applied each of the four methods (Louvain, HC, SC and k-means) on the fused similarity matrix to obtain clustering results.

Two internal validation metrics, Dunn index [45] and Connectivity [45], were employed for the first scenario to quantitatively assess the goodness of a clustering structure without relying on any clustering methods or knowing external information about class labels. The Dunn index [45] evaluates non-linear combinations of the between-group separation and the within-group compactness. The Connectivity reflects the extent of observations that are present in the same group as their neighbors in the data space. The original value of Connectivity ranges from zero to infinity, with smaller value denoting higher performance. Here we used a transform, 1/log10(Connectivity +1), to make Connectivity consistent with Dunn. The larger the score of Connectivity or Dunn, the better the separation is. The R package clValid [45] was adopted to calculate the Connectivity and Dunn index.

Additionally, we used three popular metrics to evaluate the performance of scLAPA in the context of clustering, including the ARI (Adjusted Rand Index), Jaccard and NMI (Normalized Mutual Information). The value of ARI ranges from −1 to 1, and values of NMI and Jaccard range from 0 to 1, with the higher value indicating the better performance. ARI is a widely-used metric for measuring the concordance between two clustering results. The Jaccard index quantifies the similarity between two datasets. NMI is a variation of mutual information for evaluating clustering results, which corrects the bias of the consistency caused by chance. ARI and Jaccard were calculated using the *adjustedRand* function in the R package clues [46]; NMI was obtained by the *compare* function in the R package igraph (https://igraph.org/r/).

### Bioinformatics analyses

UMAP [47] was adopted for visualization of distributions of single cells, which employs the non-linear dimensional reduction technique to group similar cells in low-dimensional space. UMAP was implemented by the *calculateUMAP* function in the scater R package [48]. For the analysis of the Arabidopsis root data, DESeq2 [49] was adopted to identify DE genes and DE poly(A) sites. First *GE-matrix* and *PA-matrix* were normalized by the median ratio method provided in DESeq2. Then the *DESeq* function was applied for DE detection. Gene or poly(A) sites with log2 fold change>=0.8 and adjusted *P*-value<=0.1 were considered as DE.

## Results

### Single-cell polyadenylation profile distinguishes cells

Recently, scRNA-seq has emerged as a unique tool to explore cell-specific gene or isoform expression in plants [50–54]. A previous study [51] utilized root-hair and nonhair cell types as models and revealed the potential of using scRNA-seq data for inferring specific cells during the process of cell-type differentiation. Here we focused on the epidermal tissue and analyzed differential expression on both gene and APA levels between root-hair and nonhair cells. A total of 294 root-hair cells and 195 nonhair cells were defined by the previous study [51]. Although both *GE-matrix* and *PA-matrix* were obtained from the same scRNA-seq data, we still found four genes exclusively present in the *PA-matrix* (Figure 1A). For example, AT1G64140, a WRKY transcription factor gene, was absent in the single-cell *GE-matrix,* while it has one poly(A) site (coord: 23803757) with much higher expression level in nonhair than in hair cells according to the *PA-matrix.* Interestingly, this poly(A) site is an annotated poly(A) site in extended 3’ UTR, which was supported by bulk 3’-seq data according to PlantAPAdb [55]. Similarly, AT3G2522, a hypothetical protein coding gene, is missing in the *GE-matrix*, while its one poly(A) site (coord: 9184927) is expressed much higher in nonhair cells than in hair cells. This poly(A) site was also annotated as a 3’ UTR site in PlantAPAdb. Moreover, 1422 genes possess at least one differentially used poly(A) site, among which 171 genes were not DE genes (Table S3). For example, AT1G59725 is a DNAJ heat shock family protein expressed in root. Although both AT1G59725 and its one poly(A) site are expressed higher in root hair cells than in nonhair cells, the difference between the two cell types characterized by the poly(A) profile is much more pronounced than that by the gene profile (Figure 1B). Further, using only the *GE-matrix*, a subset of cells are indistinguishable between hair and nonhair cell types (Figure 1C). In contrast, cells from the two cell types were clearly separated on the basis of the *PA-matrix* and two potential subpopulations of nonhair cells were observed (Figure 1C). Therefore, we anticipate that the poly(A) site expression profile may encode complementary information that is absent or insignificant in the gene expression profile, which could be useful to distinguish cell types. There is a great potential to develop integrative approaches for discerning cell identities that can properly incorporate single-cell profiling of both gene expression and polyadenylation information.

**Figure 1.**
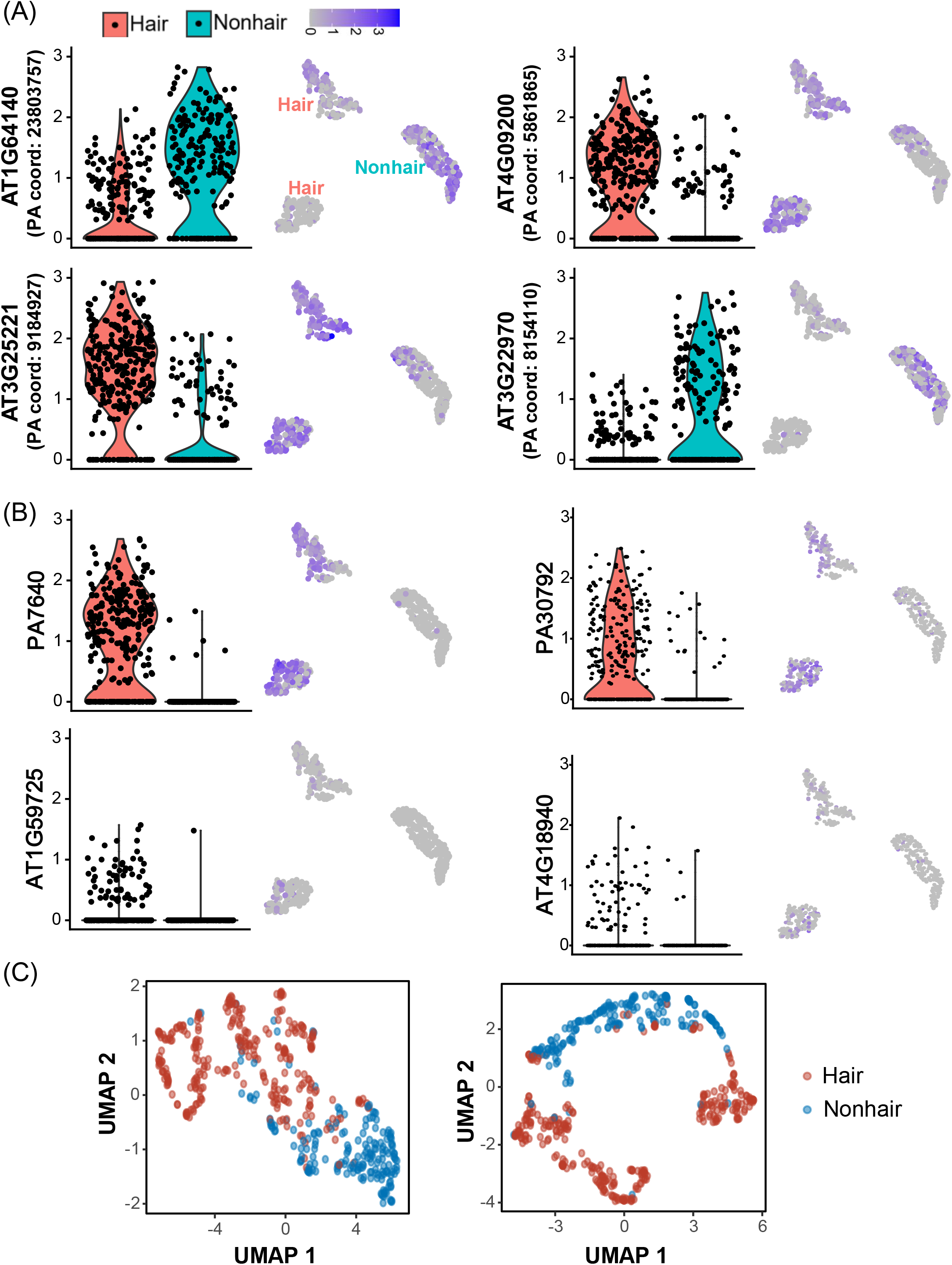
Single-cell poly(A) profile in root hair and nonhair cells. (A) Genes exclusively present in the *PA-matrix.* Four genes (AT3G22970, AT1G64140, AT4G09200 and AT3G25221) were not present in the *GE-matrix,* whereas they had at least one poly(A) site according to the *PA-matrix.* For each gene, the violin plot shows expression levels of its poly(A) site in hair and nonhair cells and the UMAP visualization shows the 2D embeddings of poly(A) profile. (B) Two example genes (AT1G59725 and AT4G18940) that are not differentially expressed (DE) but possess at least one DE poly(A) site. The upper panel places the violin plot and UMAP visualization showing the poly(A) profile of the respective gene in hair and nonhair cells. The lower panel shows the gene profile. (C) Single-cell poly(A) profile distinguishes root hair and root nonhair cells. The left plot is the UMAP representation on the basis of 171 genes that are not DE but with at least one DE poly(A) site, the right plot is the UMAP representation on the basis of poly(A) profile of the 171 genes.

### Learning cell-cell similarities with scLAPA

We proposed the scLAPA toolkit that can learn cell-cell similarities by taking advantage of the complementarities from both layers of APA isoforms and genes. Here we compared the performance of the similarity metric learned from scLAPA with other seven similarity metrics by analyzing four scRNA-seq datasets. Two metrics, Dunn and Connectivity, were adopted to quantitatively measure cell separation independent of clustering methods. Generally, scLAPA provides higher or comparable performance than other metrics across all the four datasets, whereas Pearson correlation or Euclidean has a consistently lower performance (Figures 2A and S2). In terms of both Dunn and Connectivity, scLAPA and SIMLR perform significantly better than other three metrics. Particularly, SIMLR outperforms scLAPA on the Hypothalamus data whereas scLAPA outperforms SIMLR on the Mammary data. Overall, scLAPA performs better than at least six out of the seven metrics in all the four datasets, never being the worst in any case. According to the Dunn index (Figure 2A), even for datasets where the performance of scLAPA is not the best, scLAPA is always the close match to the best. For example, the Dunn score from scLAPA on the Hypothalamus data is 0.94, which is very close to the best score (0.985 from SIMLR).

**Figure 2.**
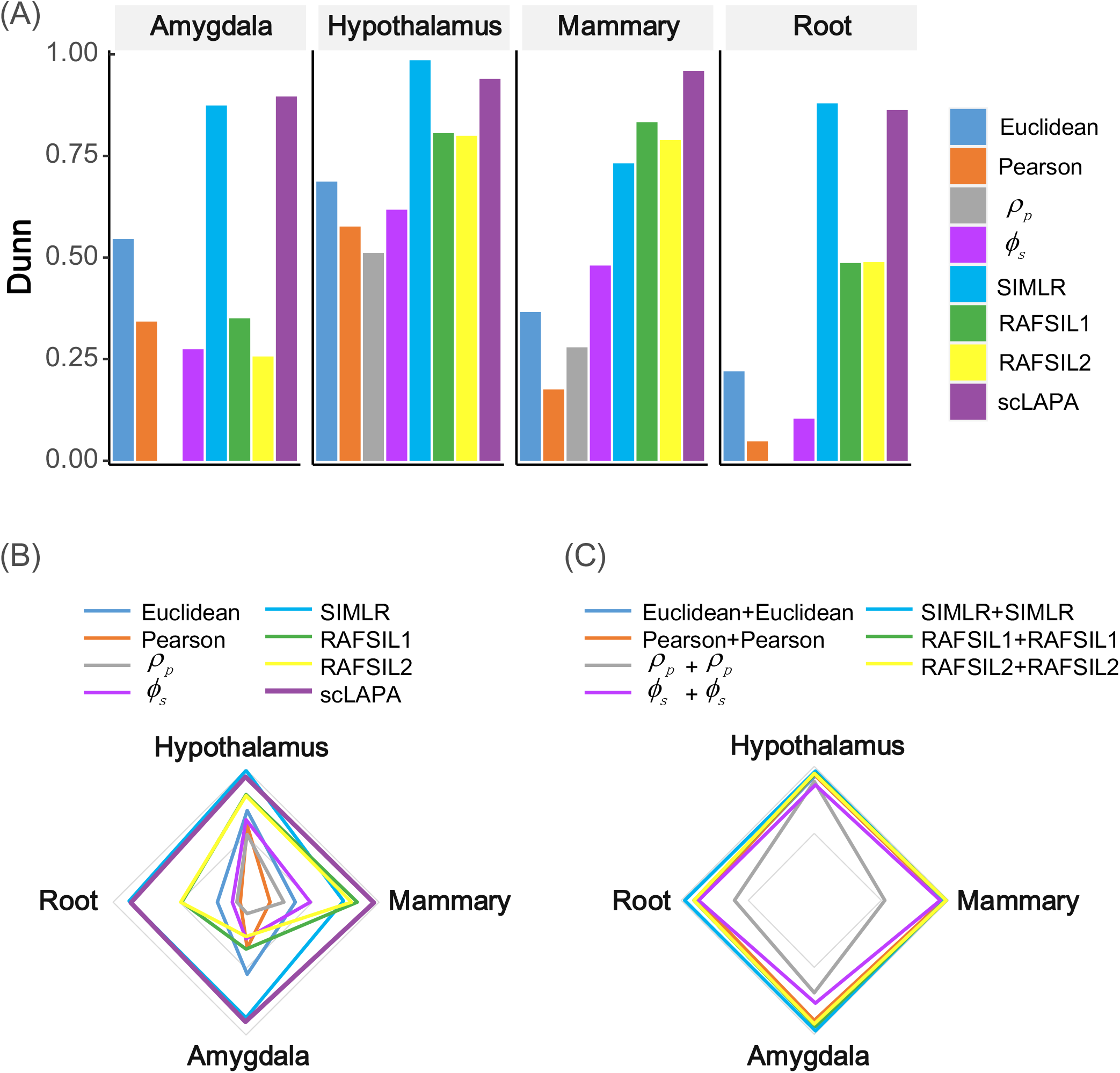
Benchmarking of similarity learning with scLAPA on four published scRNA-seq datasets. (A) The internal validation metric of Dunn was employed to measure the cell separation. (B) Radar chart showing the performance of different similarity metrics across datasets. Dataset names are shown near the vertex of the plot. Each vertex denoting the Dunn score of a metric on the respective dataset. The larger the area of a polygon displayed in a radar chart is, the higher the overall performance is. (C) Radar chart showing the performance of scLAPA with different distance metrics for distance fusion. Each vertex denotes the Dunn score of using different distance metrics on the respective dataset.

Next we used the radar chart to compare the performance of these similarity metrics more intuitively. Apparently, scLAPA and SIMLR stand out as universally better than the others, and discrepancies of performance of other six metrics across different datasets were observed (Figure 2B). For example, the overall similarity based on the RAFSIL1/2 metric is much higher on Mammary and Hypothalamus data than the other two datasets, revealing the instability of performance of RAFSIL across different datasets. In contrast, for all these four datasets, both Euclidean and Pearson correlation emerge as the worst similarity metric. In contrast, scLAPA provides a more robust result regardless of datasets. scLAPA is integrative and flexible in that different distance metrics can be chosen to learn cell-cell similarities for distance fusion. Next we examined the effect of using different distance metrics in scLAPA. The performance of scLAPA according to the Dunn index is highly robust across all datasets regardless of distance metrics used in scLAPA (Figure 2C). It is widely accepted that it is highly challenging to determine an optimal distance metric for profiling true cell-cell relationships from the complex and heterogeneous scRNA-seq data [3]. However, the integrative framework of scLAPA provides an effective solution of distance fusion by assembling results from multiple data layers into one ensemble result, which can mitigate limitations in individual similarity metrics or data layers and facilitate the generalization and adaption for different scRNA-seq datasets. Take the Hypothalamus data as an example. Apparently, the matrix with block structures obtained from scLAPA showed higher consistency with true labels than did other similarity metrics (Figure S3). Block structures learned by SIMLR are indistinguishable from background signatures; block structures learned by Pearson correlation, Euclidean or the two measures of proportion are also mixed with background signatures; block structures learned from RAFSIL are generally consistent with true structures except that cell types with small number of cells are less distinguishable. Overall, scLAPA provides more divergent clusters with higher distinction, and individual clusters obtained by scLAPA are more compact than those by other similarity metrics. These results demonstrate the ability and robustness of scLAPA in improving the cell separation across numerous scRNA-seq datasets.

### Cell type clustering with scLAPA

Cell-cell similarities learned by different similarity metrics can be adapted to other clustering methods that take similarities as inputs. Here we performed extensive comparisons of scLAPA with other seven similarity metrics by applying different clustering methods for cell clustering. First we applied Louvain [2], a graph-based method for community detection, to different similarity metrics for clustering. According to the ARI score, similarities learned by scLAPA and SIMLR significantly outperform similarities obtained from Euclidean, Pearson correlation or RAFSIL1/2 (Figure 3A). Overall, SIMLR shows similar performance with scLAPA, whereas scLAPA outperforms SIMLR in three out of the four datasets. Particularly, Euclidean and Pearson correlation present the worst performance in two datasets, Mammary and Root. Similar results were obtained in terms of other two indexes, NMI and Jaccard (Figure S4). In addition to Louvain clustering, we also investigated other three popular clustering methods, including hierarchical clustering [39], spectral clustering [40] and k-means [56], to evaluate the robustness of results by applying different clustering methods on the same similarity metric (Figures S5-7). Particularly, the performance of scLAPA and RAFSIL1/2 are robust regardless of clustering methods used, whereas scLAPA consistently outperforms RAFSIL. In contrast, SIMLR, Euclidean and Pearson correlation are very sensitive to clustering methods applied (Figure 3B). Surprisingly, although SIMLR achieves comparable performance with scLAPA based on Louvain clustering (Figure 3A), its performance is the worst using k-means or spectral clustering (Figure 3B). Take the Mammary data for example, the ARI score of SIMLR drops from 0.769 when using Louvain clustering to an extremely low median value of 0.026 when using k-means. Moreover, we noted that, ARI scores from individual runs of k-means clustering on SIMLR similarities varied greatly, revealing the relatively poor robustness of SIMLR with k-means clustering (Figure S5). These results demonstrate that the cell-cell similarity matrix learned from scLAPA is more effective and robust than competing similarity metrics in clustering cell subpopulations.

**Figure 3.**
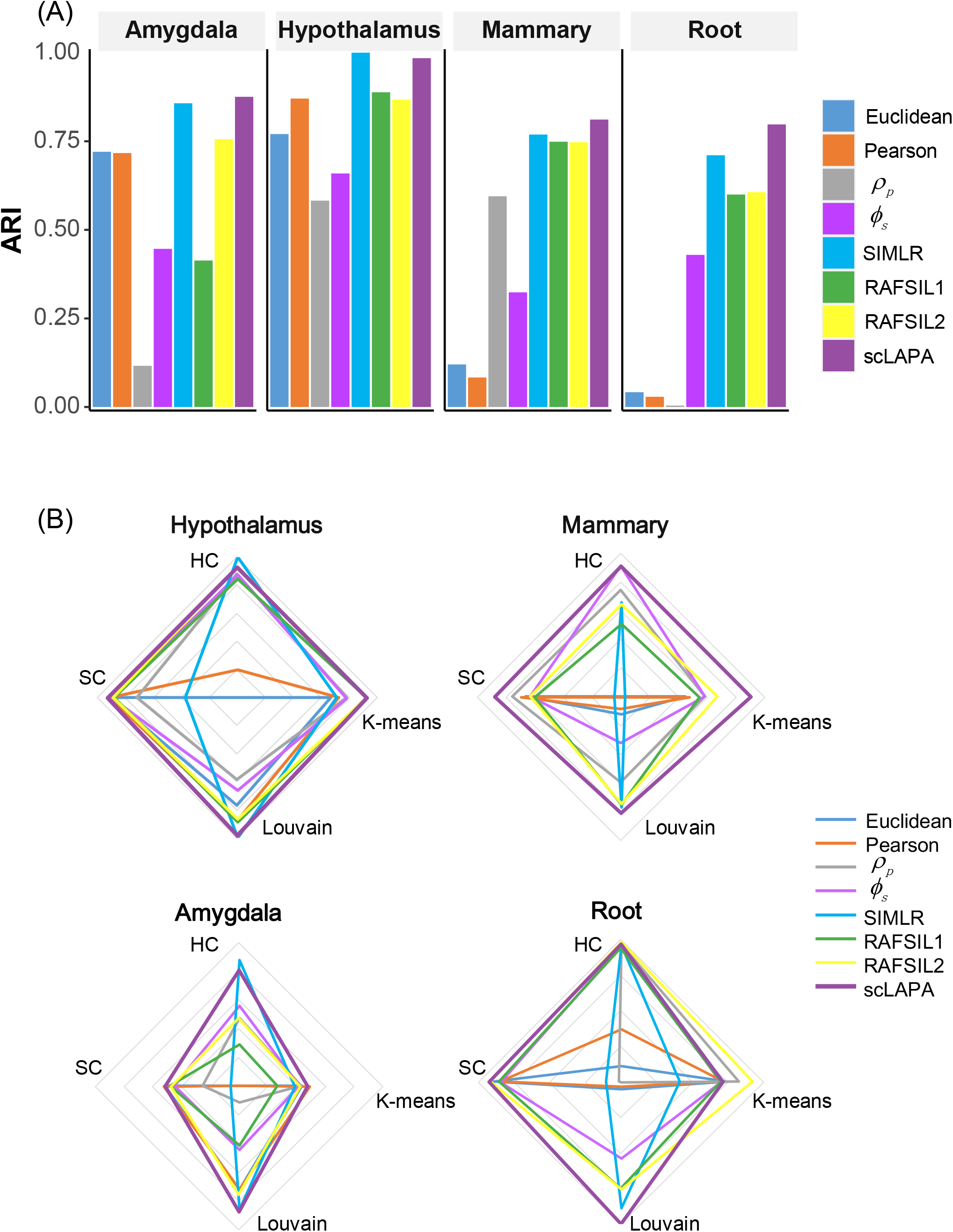
Benchmarking of similarity learning with scLAPA in the context of clustering on four published scRNA-seq datasets. (A) ARI was employed to measure the concordance between inferred and true cluster labels. Louvain clustering was applied on the similarity matrices obtained from different methods. (B) Radar charts showing ARI scores by applying different clustering methods on cell-cell similarities learned by each similarity metric. Each plot represents results of one dataset. Clustering methods are shown near the vertex of the plot. The vertex of a plot denotes the ARI score of applying a clustering method on different metrics. The larger the area of a polygon displayed in a radar chart is, the higher the overall performance is. HC, hierarchical clustering; SC, spectral clustering.

During the preparation of this manuscript, we noticed another method scDaPars [57], which quantifies and recovers APA events in single cells using standard scRNA-seq data. The authors also integrated APA information identified by scDaPars with imputed gene expression by similarity network fusion to reveal novel cell subpopulations during human embryonic development. Different from scDaPars that employs the (imputed) percentage of distal poly(A) site usage index (PDUI) to measure APA usage, scLAPA directly utilizes raw poly(A) expression profile. Here we compared the performance of scLAPA and scDaPars by applying them to the four scRNA-seq datasets in our benchmarking analysis. Following the process in Gao et al. [57], we calculated PDUI based on the *PA-matrix* and imputed APA profiles using scDaPars. Then we applied five similarity metrics on the scDaPars-imputed APA profile and the *GE-matrix* to generate *scDaPars-dist* and *GE-dist*, respectively. After fusing the two distance matrices with SNF, we applied Louvain clustering on the fused cell-cell similarities to cluster cells. According to the ARI score (Figure 4), scLAPA significantly outperforms scDaPars on all the four datasets. Particularly, ARI scores of scDaPars with different similarity metrics varied greatly whereas the performance of scLAPA is robust with different similarity metrics (Figure 4 vs. Figure 2C), revealing that the poly(A) expression profile used in scLAPA is more efficient and robust than the PDUI profile used in scDaPars for clustering cells.

**Figure 4.**
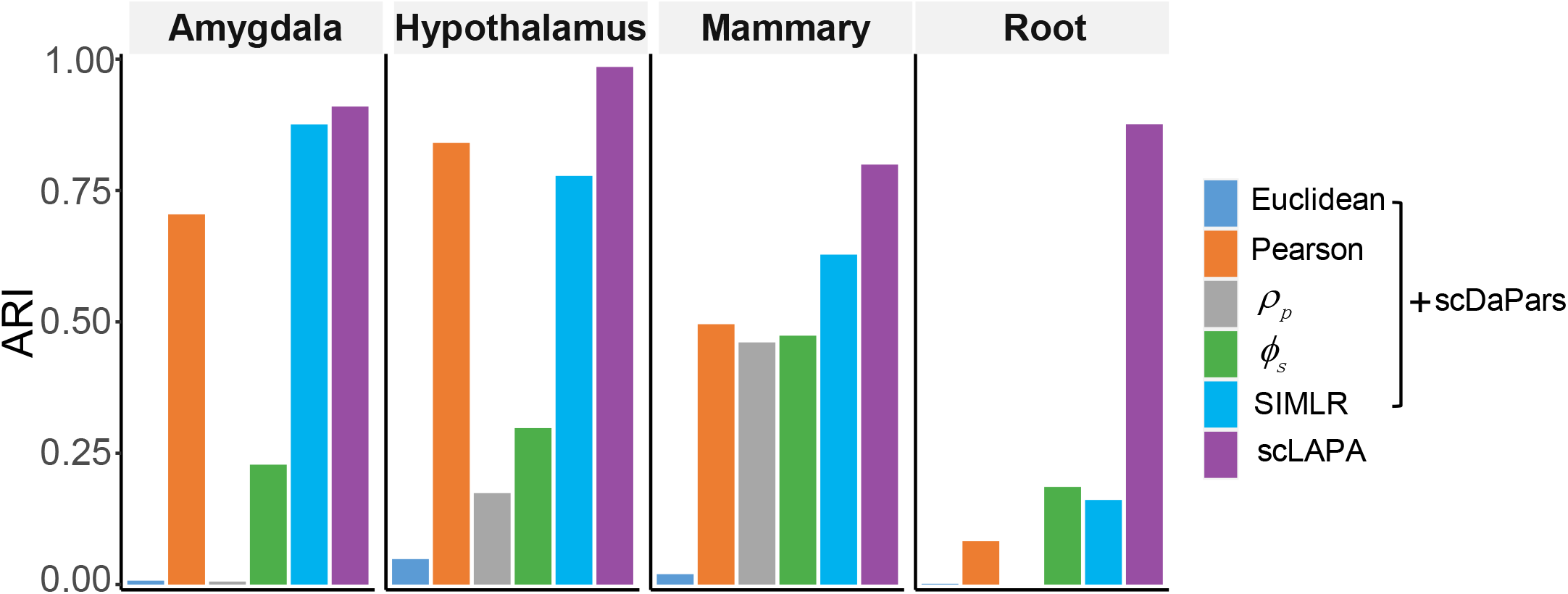
Comparison of performance between scLAPA and scDaPars across four scRNA-seq datasets. Five similarity metrics were applied on the scDaPars-imputed PDUI profile and the *GE-matrix* to generate *scDaPars-dist* and *GE-dist*, respectively. After fusing the two distance matrices with SNF, Louvain clustering was applied on the fused cell-cell similarities to cluster cells. We did not include RAFSIL in this experiment due to its slow calculation speed. For scLAPA, Pearson correlation was used for similarity learning and Louvain was used for clustering.

Next we expanded the benchmarking analysis by comparing clustering results of scLAPA with other single-cell clustering methods that directly take the gene-cell expression matrix as input without an explicit procedure of similarity learning. Specifically, we included five popular tools for comparison, including SC3 [6], Seurat v3 [13], SINCERA [42], SNN-Cliq [43] and dynamicTreeCut [44]. According to the ARI score, scLAPA achieves generally higher or comparable performance than other methods, whereas dynamicTreeCut provides a consistently lower performance (Figure 5). Similar results were observed using indexes of Jaccard or NMI (Figure S8). Specifically, scLAPA provides the best ARI score in three out of the four datasets (Figure 5). For the Hypothalamus data where SC3 performs the best, scLAPA presents very close ARI score to SC3 (scLAPA=0.985; SC3=0.99). Particularly, for three datasets (Mammary, Hypothalamus and Root), ARI scores of individual SC3 runs varied greatly, reflecting the performance of SC3 may be unstable on some kinds of datasets. Overall, the performance of scLAPA is robust and consistently high across diverse scRNA-seq datasets.

**Figure 5.**
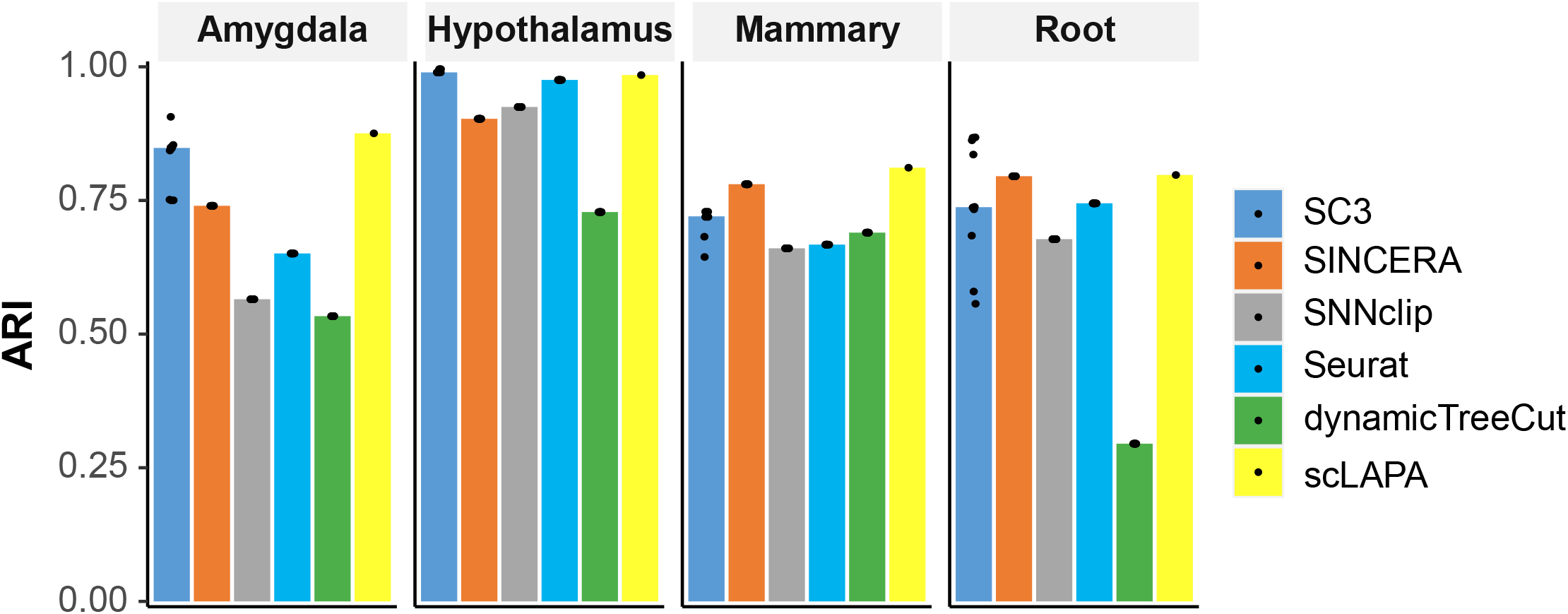
ARI scores from six clustering methods across four scRNA-seq datasets. For scLAPA, Pearson correlation was used for similarity learning and Louvain was used for clustering.

### scLAPA identifies hidden subpopulations of cells

We next applied scLAPA on the human PBMC 4k dataset from 10x Genomics for cell type clustering. First we examined the cell type composition of the PBMCs by applying Seurat to the gene-cell expression matrix (*GE-matrix*). Ten distinct cell clusters were yielded (Figure 6A). Based on the expression of known markers (Table S2), nine clusters were annotated. Up to 13,512 poly(A) sites from 9601 genes were identified from the raw RNA-seq data with scAPAtrap. We learned cell-cell similarities with scLAPA by jointly considering expression profiles of APA isoforms and genes. After applying Louvain clustering on the cell-cell similarity matrix, 14 cell clusters were obtained and 11 clusters were successfully annotated. These 11 clusters covered the nine clusters identified by Seurat and contained two new small clusters (Figure 6B). Both subclusters were supported by the expression patterns of markers, suggesting that they represented distinct cell types. One subcluster was annotated as regulatory T cell on the basis of elevated expression of three markers, *CCR10, FOXP3* and *IL2RA* (Figure S9). Depending only on the gene expression profile, regulatory T cells were not well resolved and are indistinguishable among other T cells (Figure 6A). Although the gene expression of the marker *CCR10* is sparse and weak among T cells, we could still distinguish clearly regulatory T cells from other T cell types according to the UMAP visualization of the gene expression profile (Figure 6C). Particularly, *CCR10* has four annotated poly(A) sites according to APASdb [58], whereas only one poly(A) site was identified from scRNA-seq data. This is not unexpected as the bulk 3 ‘-seq data contain more diverse tissue samples than the PBMC data and scRNA-seq data is generally too sparse to identify all poly(A) sites. However, we have shown that, even for a single poly(A) site, it could encapsulate useful information beyond the gene expression profile (Figure 1). The other subcluster where cell markers such as *PPBP* and *PF4* are expressed, was annotated as megakanyocyte progenitors (Figures 6D and S10). According to the *PA-matrix, PPBP* carries three poly(A) sites, and five poly(A) sites of *PPBP* were annotated in APASdb. These three poly(A) sites were all highly expressed in megakanyocyte progenitors (cluster 10) (Figure 6E). These results demonstrate that scLAPA facilitates the capture and identification of hidden subpopulations of cells that are unrecognizable based on the gene expression profile alone.

**Figure 6.**
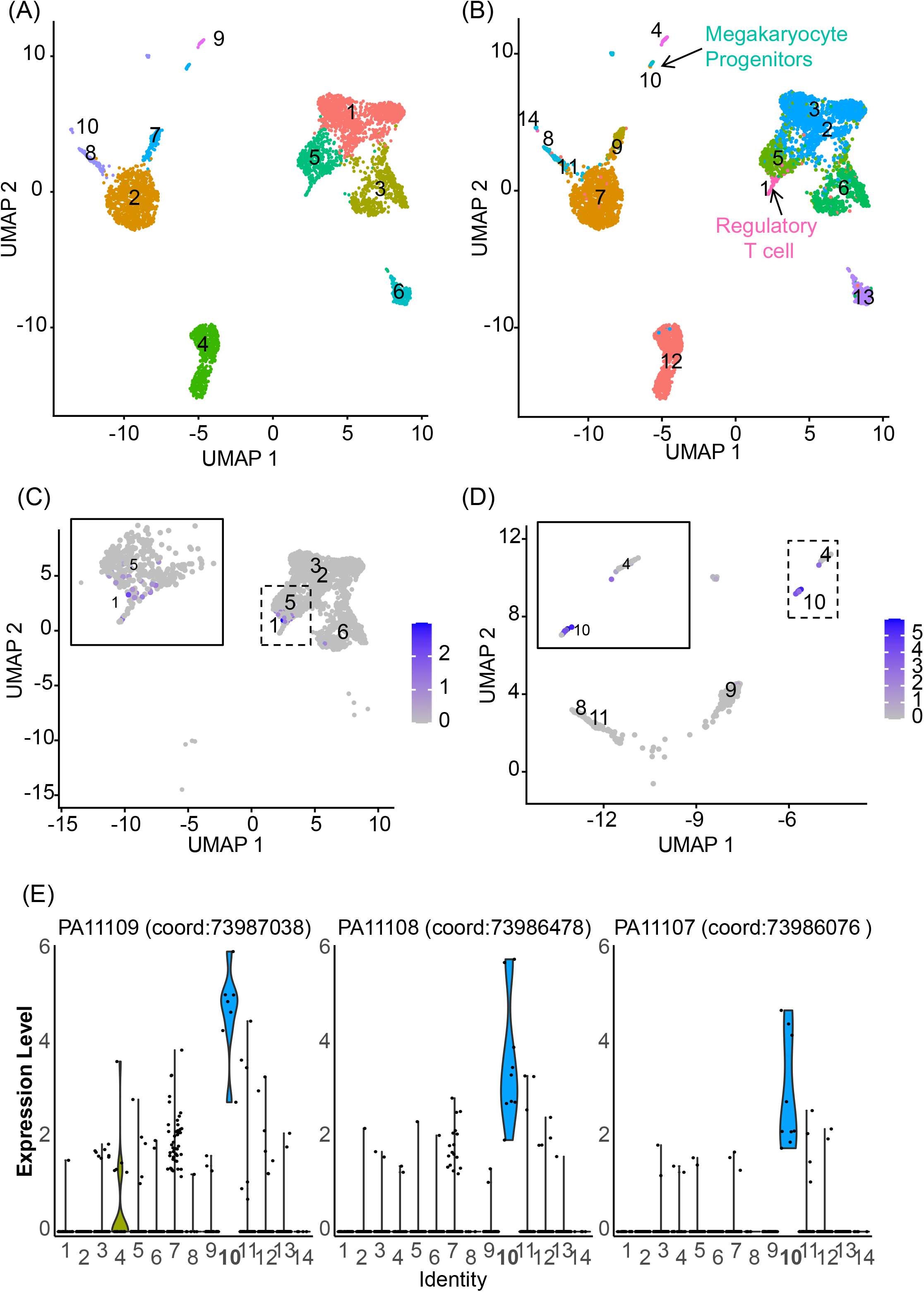
scLAPA identifies hidden subpopulations of cells from human PBMCs. (A) UMAP representation of Seurat’s clustering results on the basis of *GE-matrix.* Ten clusters were obtained and nine were annotated with known cell types: Naive T cell (1), CD14+ Monocytes (2), CD8+ T cell (3), B cell (4), CD4+ Memory T (5), NK cell (6), CD16+ Monocytes (7), Monocyte Derived Dendritic (8, 10) and Plasmacytorid Dendritic (9). (B) UMAP representation of scLAPA’s clustering results on the basis of *GE-matrix* and *PA-matrix.* Fourteen clusters were obtained and 11 clusters were annotated with known cell types: Regulatory T cell (1), Naive T cell (2, 3), Plasmacytorid Dendritic (4), CD4+ Memory T (5), CD8+ T cell (6), CD14+ Monocytes (7), Monocyte Derived Dendritic (8, 11, 14), CD16+ Monocytes (9), Megakaryocyte Progenitors (10), B cell (12) and NK cell (13). The two arrows mark two new subpopulations of cells identified by scLAPA. (C) Gene expression of *CCR10* distinguishes regulatory T cells from other T cell types according to the UMAP visualization of the gene expression profile. The details in the dashed line box are shown in the solid line box. (D) Gene expression of *PPBP* distinguishes megakanyocyte progenitors from other cell types. (E) Three poly(A) sites of *PPBP* are all highly expressed in megakanyocyte progenitors.

## Discussion

scLAPA is an integrative framework for learning association for single-cell transcriptomics by leveraging expression profiles of genes and APA isoforms in individual cells, which highlights the inclusion of polyadenylation signatures for improving cell type clustering and discovering new cell types. The effectiveness of scLAPA for cell-cell similarity learning and cell type clustering is evidenced by comparisons with various similarity metrics and single-cell clustering methods on several scRNA-seq datasets. scLAPA has a number of desirable features. First, scLAPA incorporates existing tools to extract and quantify poly(A) sites directly from scRNA-seq, which augments the gene-level analysis with additional layer of APA information without altering the scRNA-seq protocol or performing additional sequencing experiment. Second, by employing the strategy of similarity network fusion, scLAPA jointly considers expression profiles at both levels of APA isoforms and genes for learning highly informative cell-cell similarities. Third, in contrast to many other methods that cluster cells without explicit similarity learning step, scLAPA provides two independent but connected modules for similarity learning and cell clustering, each with various methods for users’ choice. Accordingly, users can freely combine different similarity metrics and clustering methods in scLAPA to evaluate the clustering results for any given dataset. Fourth, the framework of scLAPA is highly flexible, which can be seamlessly embedded into most existing scRNA-seq pipelines or tools for downstream analyses, such as dimension reduction, cell type clustering and differential expression analysis. Accordingly, existing tools, such as those designed for dropout imputation, normalization and similarity learning, can also be easily incorporated into scLAPA.

With scLAPA, distinct cell-cell similarity networks can be effectively learned from profiles of gene expression and polyadenylation separately by various similarity metrics. scLAPA then employed the strategy of similarity network fusion for scalable and robust integration of similarity networks learned from different data layers. This strategy has the advantage to exploit complementarities in distinct data layers for fully profiling the spectrum of underlying data. Moreover, the consensus set of cell-cell interactions and associations from the APA layer and the gene layer can be learned from the given data, mitigating noise and dropouts in conventional gene-cell expression profile and thus enhancing accuracy for downstream analyses. By combining expression profiles of APA and gene through similarity network fusion, we found two hidden subpopulations of PBMCs that were undetectable using only gene expression data (Figure 6). Moreover, the augmentation of gene expression profiles with polyadenylation information enhances single-cell clustering results and generates more discriminative cell types (Figures 2–5). As a comprehensive toolkit, scLAPA provides a unique strategy to improve cell type clustering and discover novel cell types, by combining gene expression with polyadenylation information at single-cell resolution.

scLAPA consists of three core function modules, including learning cell-cell similarities, distance fusion and clustering. Currently, numerous methods are available to learn cell-cell similarities or cluster cells with reasonable accuracy [3]. However each method has its own strengths and limitations, and it is extremely challenging, if not impossible, to determine an optimal method for all kinds of datasets as different methods may exploit different characteristics in the data [59]. Moreover, some similarity metrics may be overly dependent on downstream clustering methods, exacerbating difficulties in choosing a universally applicable combination of similarity and clustering methods. For example, based on the *GE-matrix* alone, similarities learned from SIMLR provide an overall high performance across datasets in terms of internal validation indexes (Figure 2A). However, SIMLR is highly dependent on downstream clustering methods for single-cell clustering; it achieves high performance with Louvain clustering (Figure 3A), whereas its performance drops sharply with k-means or spectral clustering (Figure 3B). In contrast, our benchmarking analyses showed that performances of scLAPA are robust and consistently high across diverse datasets regardless of distance metrics or clustering methods selected in scLAPA (Figures 2–5). The unique strength of scLAPA may be due to that it efficiently fuses rich structures stored in *GE-matrix* as well as the accompanied *PA-matrix,* thus can amplify biological signals and augment cell-cell relationships.

scLAPA is an easy-to-use and highly flexible framework. The input of scLAPA is the *GE-* and *PA-matrix,* without using any priori biological information. Even with raw scRNA-seq data, it is easy obtain the prerequisite *GE-matrix* and/or *PA-matrix* using various tools, e.g. Cell Ranger for *GE-matrix,* scAPAtrap and Sierra for *PA-matrix.* Lately another tool, scDaPars [57], was proposed to quantify and recover APA usages from scRNA-seq data, which uses the relative usage of the distal poly(A) site called PDUI to measure a gene’s APA usage. With scDaPars, Gao et al. [57] analyzed cell-type-specific APA regulation and discovered hidden cell subpopulations from cancer and human endoderm differentiation scRNA-seq data. In scLAPA the input *PA-matrix* can be replaced with any other gene-cell-like matrix, thus the scDaPars-imputed PDUI matrix can be used readily in scLAPA for downstream cell type clustering. However, although the scDaPars-imputed PDUI profile seems to be effective in revealing APA dynamics among cell types in the previous study [57], we found that, for cell type clustering, the performance with the *PDUI-matrix* is much lower and less robust than that with scLAPA’s *PA-matrix* (Figure 4). This may be due to several reasons. First, only genes with at least two 3’ UTR poly(A) sites can be used for scDaPars’ PDUI calculation, consequently the *PDUI-matrix* is much more sparse than the *PA-matrix* and information encoded in genes with single poly(A) site is lost. Second, although the PDUI profile can be imputed with scDaPars, limited information in the highly sparse *PDUI-matrix* confounds reliable imputation and may lead to propagation of errors or noises during the imputation process. Third, unlike scLAPA which is specifically designed for learning cell-cell similarities and cell type clustering, the main function of scDarpas is to analyze cell-type-specific APA dynamics and identify novel APA-related cell types. We anticipate that the *PA-matrix* used in scLAPA may contain more comprehensive and reliable information than the *PDUI-matrix* or the imputed *PDUI-matrix*, which can significantly enhance the accuracy of cell type clustering. Overall, the *PA-matrix* is simple but effective which can be easily obtained from scRNA-seq data by various tools, making it more convenient to use scLAPA for scRNA-seq analyses.

For practical application purpose, the current version of scLAPA implements seven similarity metrics and four clustering methods for users’ choice, which allows users to investigate their own strategies for evaluation of the effect of different combinations of distance metrics and clustering methods. Moreover, scLAPA is easily expandable in that additional distance metrics or clustering methods can be readily incorporated. Meanwhile, scRNA-seq preprocessing steps, such as dropout imputation and normalization, can also be easily applied before similarity learning. scLAPA can also be used as a plug-in architecture for most existing scRNA-seq pipelines for similarity learning and cell clustering.

### Key Points

- We proposed a computational toolkit called scLAPA for learning association for single-cell transcriptomics from scRNA-seq data.
- scLAPA improves cell-cell similarity learning and cell type clustering by integrating single-cell profiling of gene expression and alternative polyadenylation.
- Objective benchmarking analyses using diverse scRNA-seq datasets demonstrate higher performance and robustness of scLAPA than competing methods in cell-cell similarity learning and cell type clustering.
- scLAPA discovers hidden subpopulations of cells that are unrecognizable based on the gene expression profile alone.

## Supplementary Data

The file of supplemental materials contains all the Supplementary Figures, and Tables.

