## Supplemental Figures for "Learning association for single-cell transcriptomics by integrating profiling of gene expression and alternative polyadenylation"

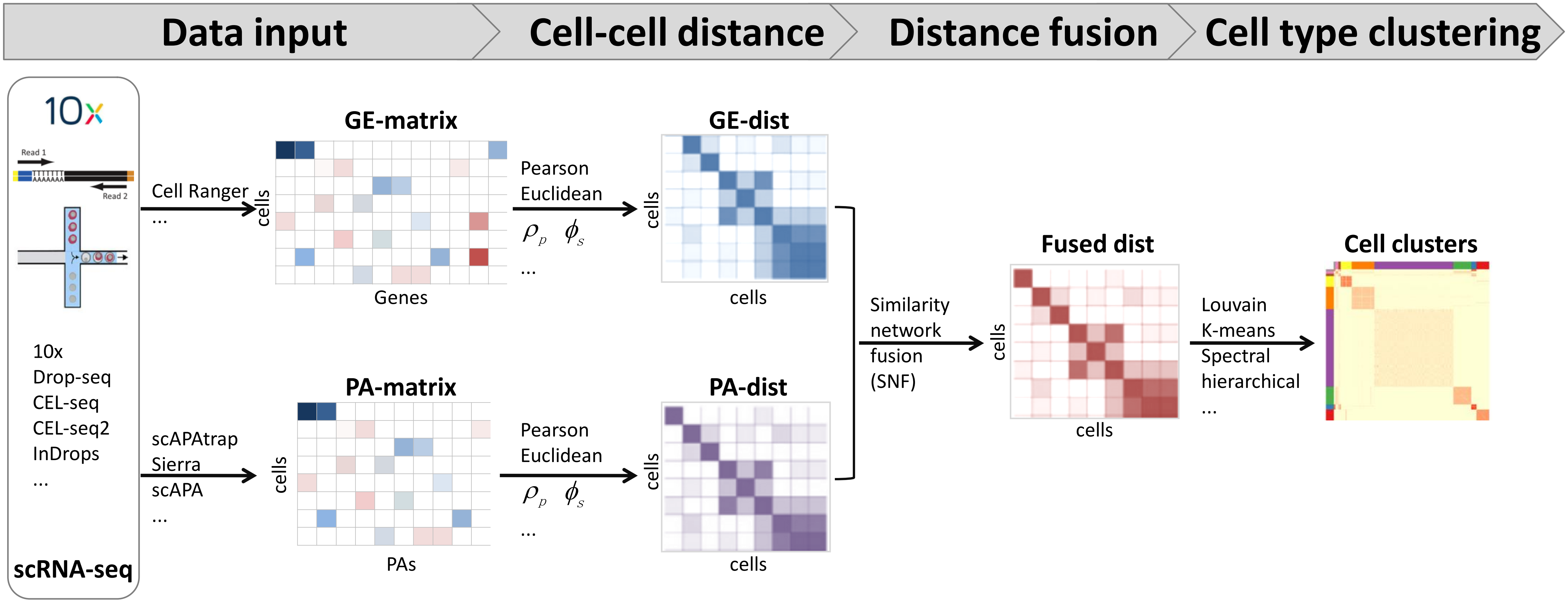


**Figure S1. Schema of scLAPA.** scLAPA consists of four modules: (i) the input module, (ii) cell-cell distance, (iii) distance fusion, (iv) cell type clustering. GE, gene expression; PA, poly(A) site.

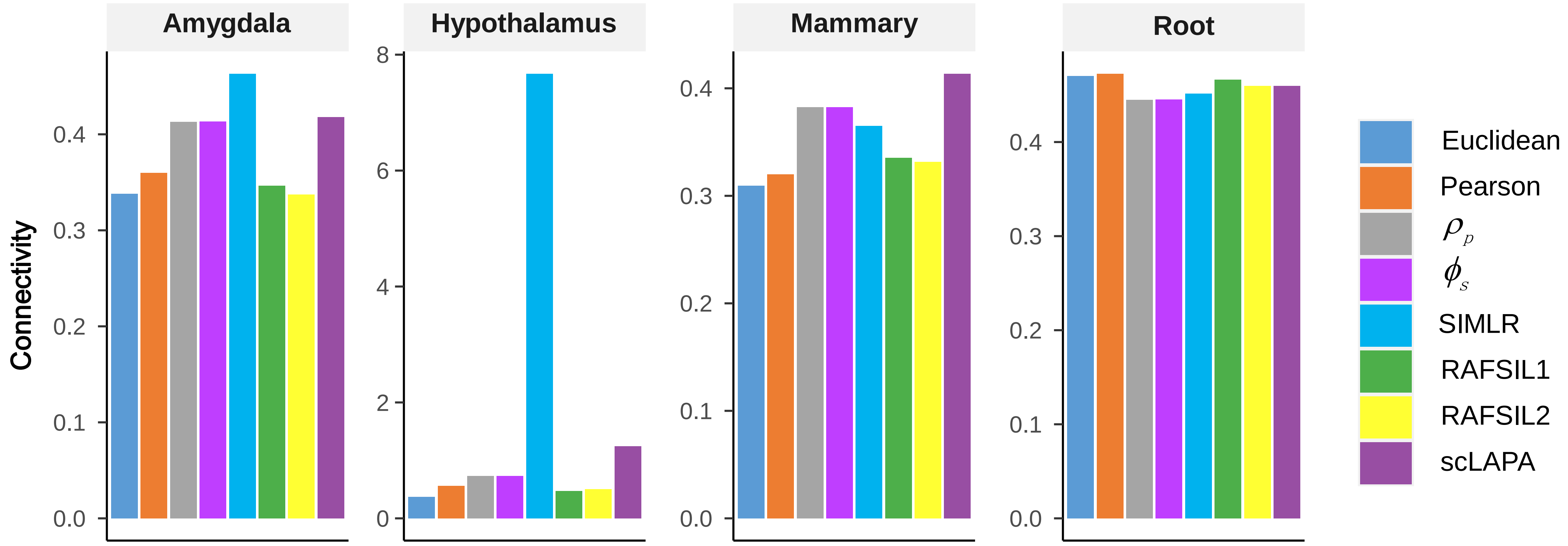


**Figure S2. Benchmarking of similarity learning with scLAPA on four published scRNA-seq datasets.** The internal validation metric of Connectivity is employed to measure the cell separation.

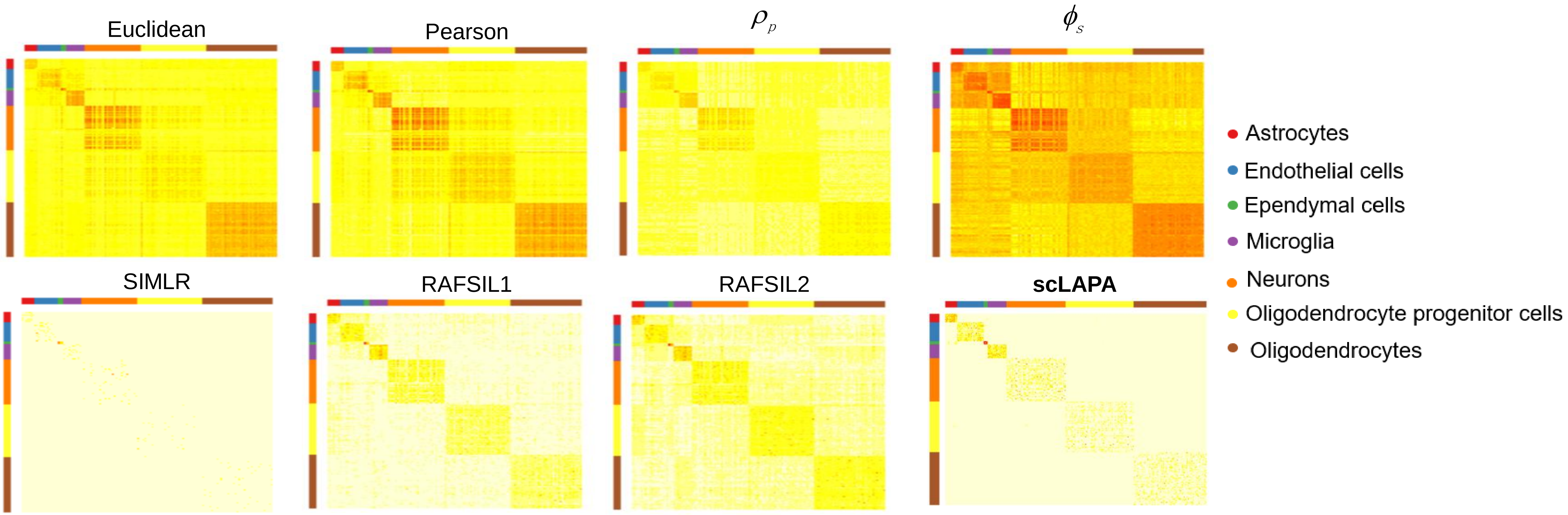


**Figure S3. Heatmap for similarities learned from the Hypothalamus data by different similarity metrics.**

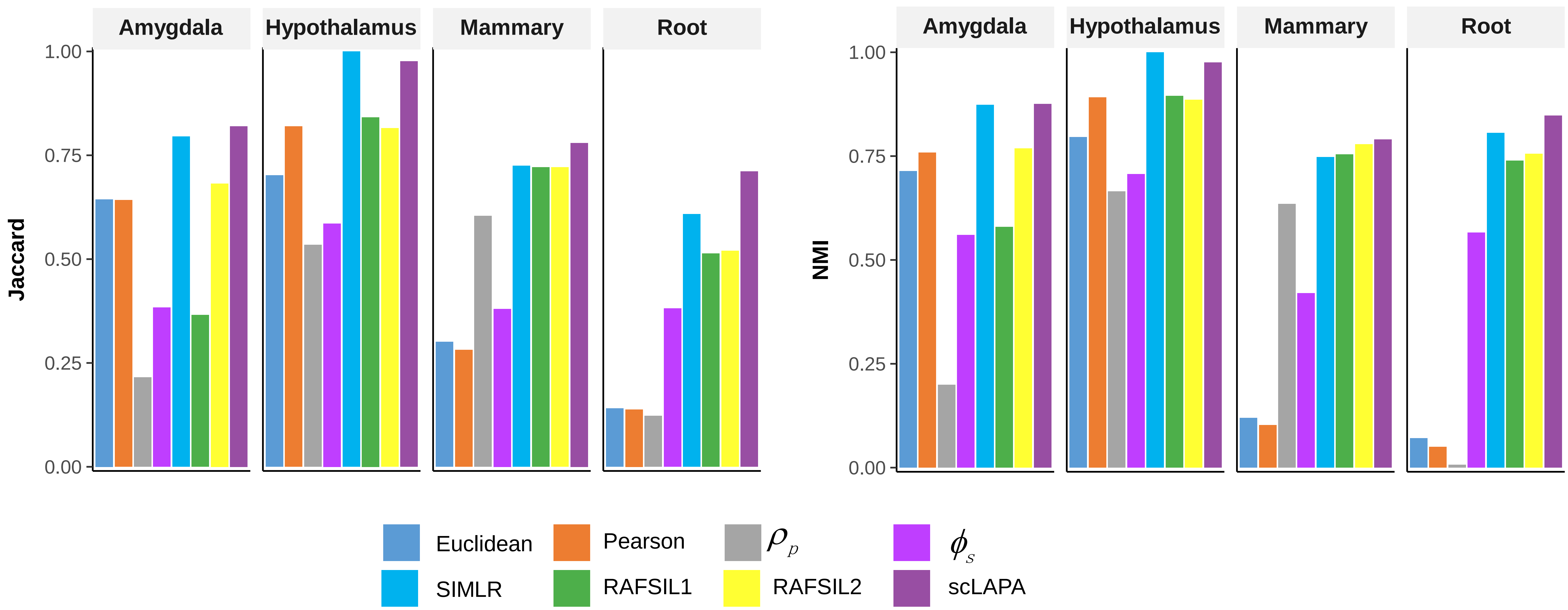


**Figure S4. Benchmarking of similarity learning with scLAPA in the context of clustering on four published scRNA-seq datasets.** (A) Jaccard is employed to measure the concordance between inferred and true cluster labels. Louvain clustering is applied on the similarity matrices obtained from different methods. (B) As in B, except that NMI score is calculated.

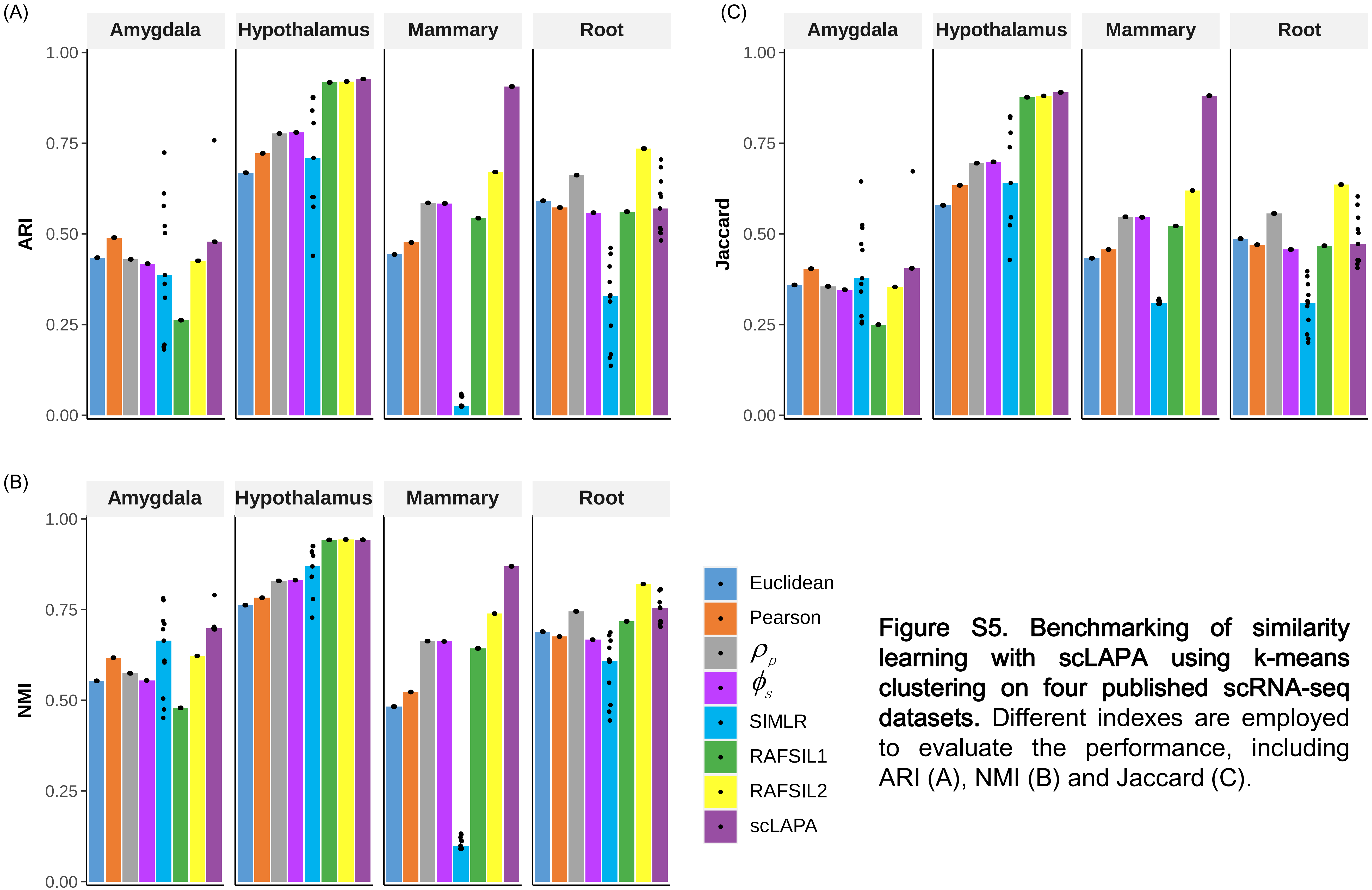


**Figure S5. Benchmarking of similarity learning with scLAPA using k-means clustering on four published scRNA-seq datasets.** Different indexes are employed to evaluate the performance, including ARI (A), NMI (B) and Jaccard (C).

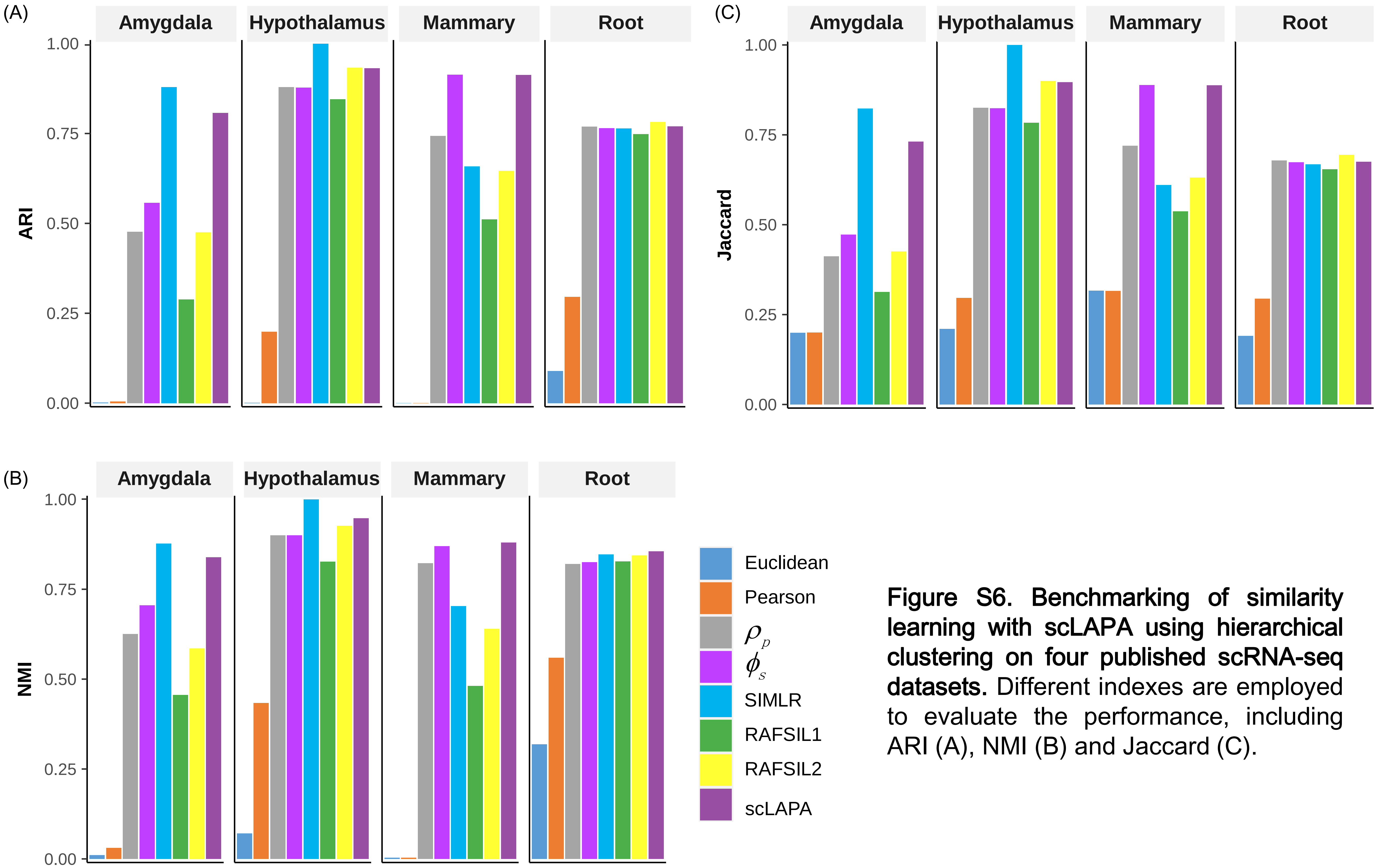


**Figure S6. Benchmarking of similarity learning with scLAPA using hierarchical clustering on four published scRNA-seq datasets.** Different indexes are employed to evaluate the performance, including ARI (A), NMI (B) and Jaccard (C).

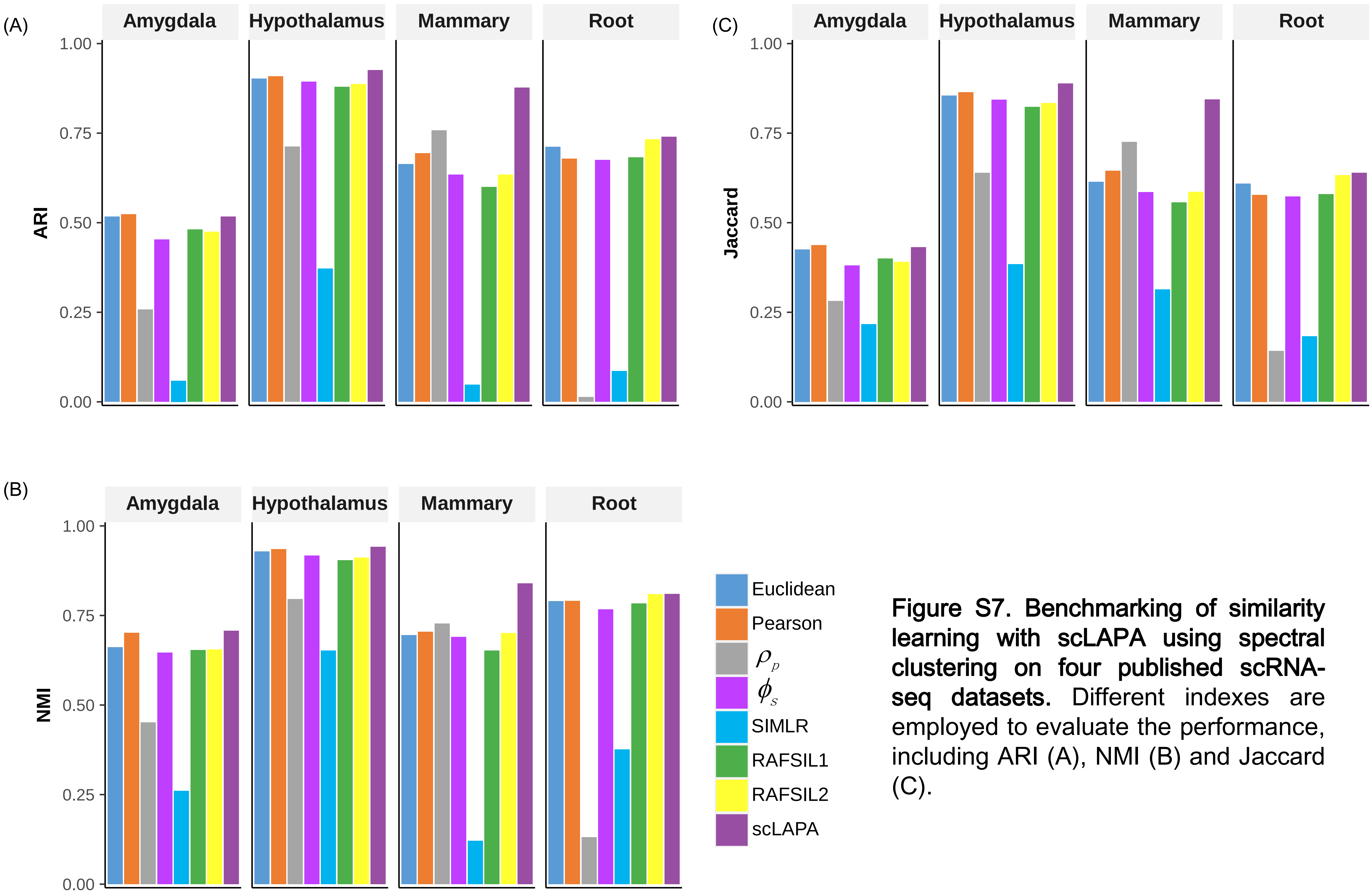


**Figure S7. Benchmarking of similarity learning with scLAPA using spectral clustering on four published scRNA-seq datasets.** Different indexes are employed to evaluate the performance, including ARI (A), NMI (B) and Jaccard (C).

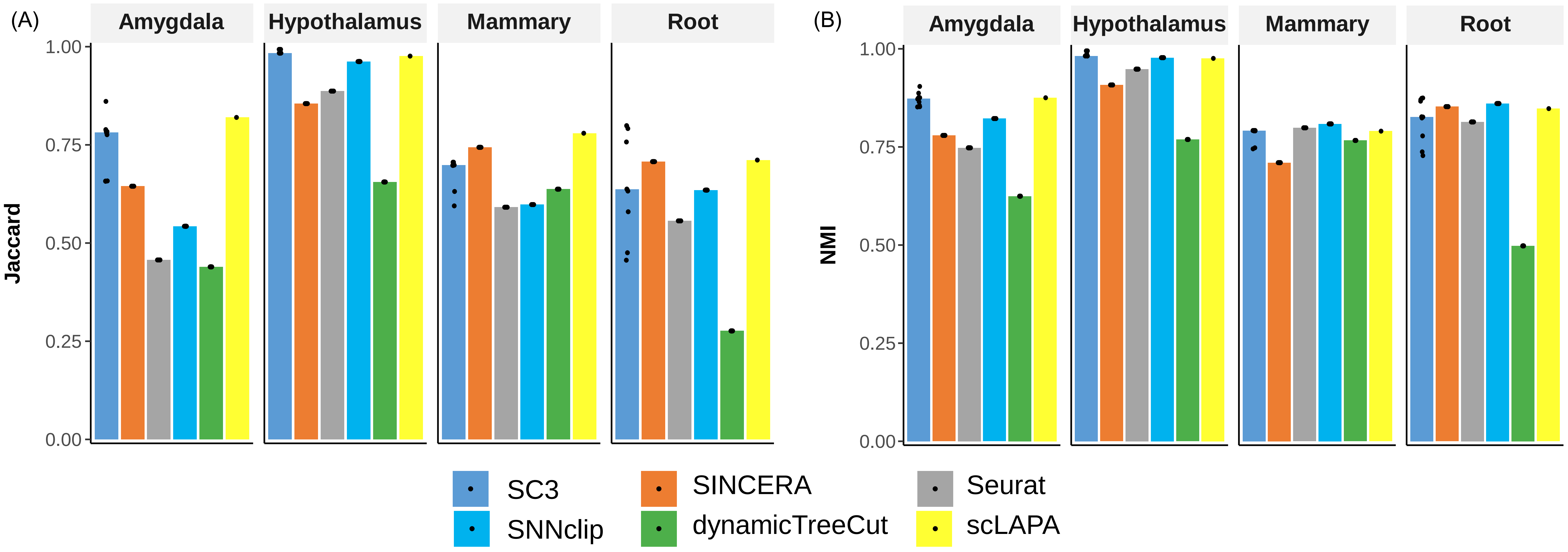


**Figure S8. Benchmarking of scLAPA on single cell clustering across four scRNA-seq datasets.** Different indexes are employed to evaluate the performance, including Jaccard (A) and NMI (B).

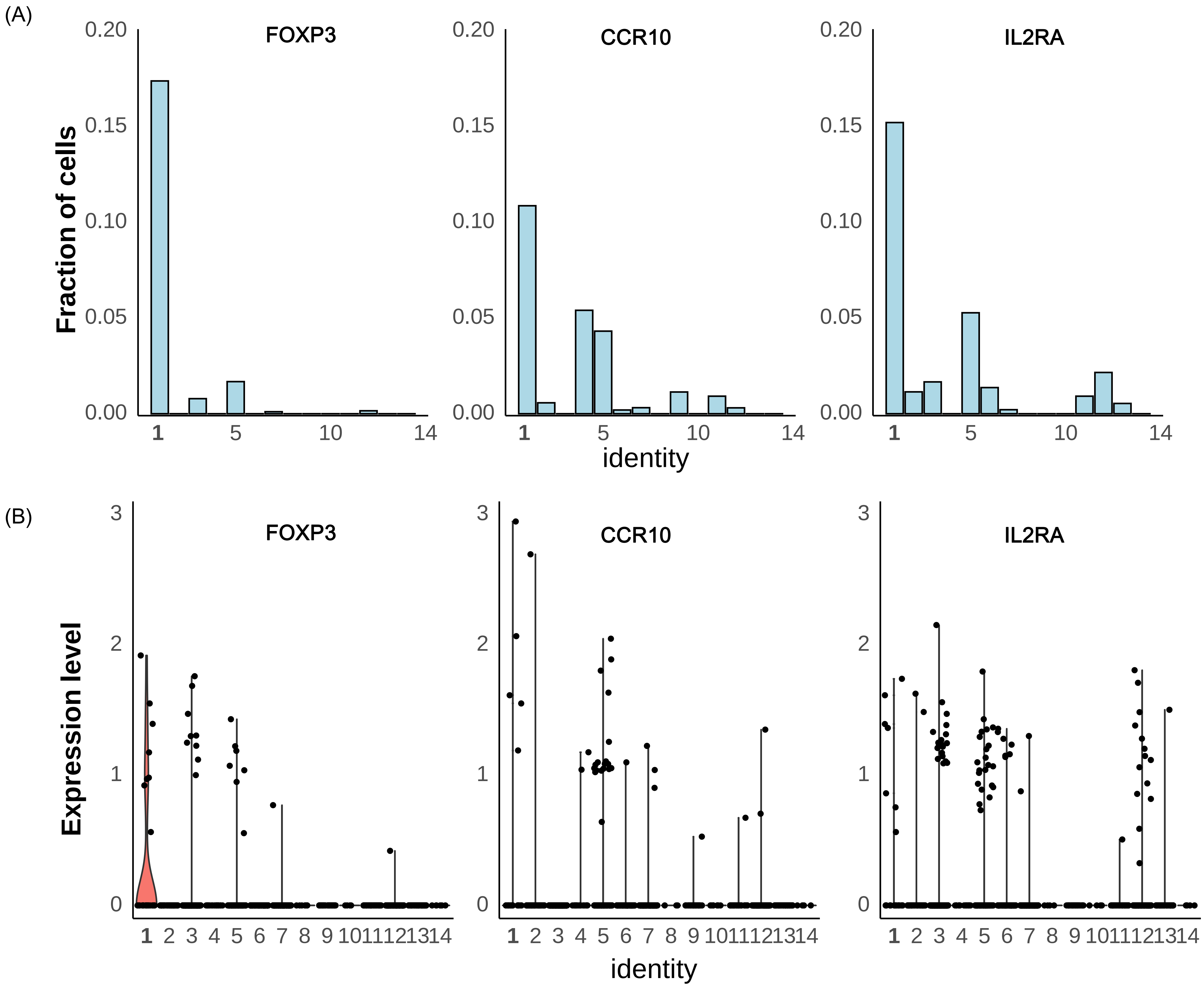


**Figure S9. Three markers (CCR10, FOXP3 and IL2RA) of regulatory T cell (cluster 1).** (A) Fraction of cells where the marker gene is expressed. (B) Expression levels of the marker gene in each cell cluster. The x-axis denotes the cluster id.

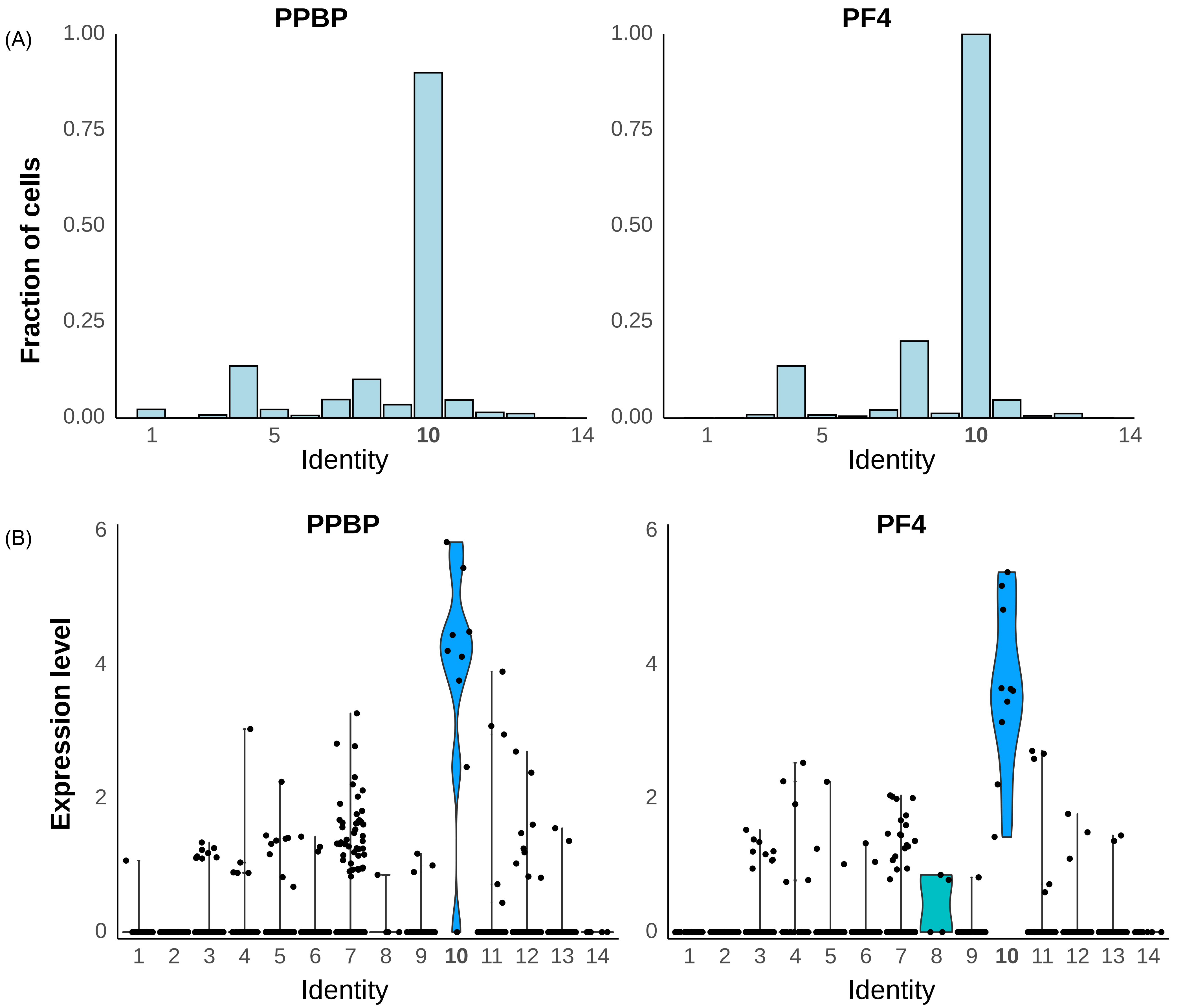


**Figure S10. Two markers (PPBP and PF4) of megakaryocyte progenitors (cluster 10).** (A) Fraction of cells where the marker gene is expressed. (B) Expression levels of the marker gene in each cell cluster. The x-axis denotes the cluster id.
